# Naltrexone has variable and schedule-dependent effects on oral squamous cell carcinoma cells

**DOI:** 10.1101/2025.08.13.670157

**Authors:** Sahar Kazmi, Erica Sanford, Zaid A. Rammaha, Feng Gao, Linda Sangalli, Cai M. Roberts

**Affiliations:** College of Dental Medicine - Illinois, Midwestern University, Downers Grove, IL, USA; Biomedical Sciences Program, Midwestern University, Downers Grove, IL, USA; Department of Pharmacology, Midwestern University, Downers Grove, IL, USA

**Keywords:** Oral squamous cell carcinoma, naltrexone, opioid growth factor receptor, chemotherapy

## Abstract

Oral squamous cell carcinoma is marked by profound differences in survival between localized and disseminated disease. While at its site of origin, oral cancer is frequently treated with surgery and/or radiation, leading to approximately 70% five-year survival. In metastatic cases, chemotherapy is preferred, but drug resistance and therapy failure is common in this setting, leading to five-year survival below 40%. New therapeutic approaches are therefore needed to supplement existing chemotherapies and improve outcomes. Naltrexone, an opioid antagonist most well-known for treatment of substance abuse, has been evaluated at low doses in other contexts, including inflammation, autoimmunity, and other forms of cancer. In this study, we sought to determine if intermittent dosing of naltrexone would impact oral cancer cell survival, either as a single agent or in combination with traditional chemotherapy. We found that its effects alone were negligible, but that combination with cytotoxic drugs may have promise as a novel treatment strategy. Further study and optimization will be needed to determine if these findings will be of clinical benefit.

## 1. Introduction

Oral Squamous Cell Carcinoma (OSCC) is a major public health burden and the most common malignancy affecting the oral cavity (Chamoli et al., 2021). It ranks among the top 20 most common cancers worldwide and accounts for approximately 220,000 new cases globally (Jiang^*^, Wu^*^, Wang, & Huang, 2019). The disease trajectory and survival outcome vary profoundly depending on the stage of diagnosis. When OSCC is detected early and remains confined to the primary site, standard treatment typically consists of surgical resection with or without adjuvant radiation (Montero & Patel, 2015), resulting in five-year survival rates of about 70% (Zanoni et al., 2019). In contrast, metastatic OSCC often necessitates systemic chemotherapy, where treatment efficacy is hindered by drug resistance, side effects, and limited durability of the response, contributing to a five-year survival rate of less than 40% (Beckham et al., 2019; Y. Cheng, Li, Gao, Zhi, & Ren, 2021; Minhas, Kashif, Altaf, Afzal, & Nagi, 2017; Shah, Shah, Thakkar, & Parikh, 2023; Wang, Zhang, & Chen, 2019). Alarmingly, 60% of OSCC cases are identified at an advanced stage (stage III or higher), dictating a much poorer prognosis (Seoane-Romero et al., 2012). Despite advances in surgical techniques, radiation protocols, and chemotherapeutic agents, the overall five-year survival rate has plateaued at around 50% for the past two decades (Warnakulasuriya, 2009). Further, projections from the Global Cancer Observatory indicate that by 2040, the incidence rate of OSCC will rise by approximately 40%, and its associated mortality will increase by 30%, underscoring the urgency for improved therapeutic approaches (Barsouk, Aluru, Rawla, Saginala, & Barsouk, 2023; Tan et al., 2023).

Recent studies have investigated novel agents to supplement traditional management options. Naltrexone (NTX) is a non-selective competitive opioid receptor antagonist approved by the Food and Drug Administration (FDA) for addiction treatment at dosages between 50 and 150 mg. At low-dose (1-5 mg), NTX has been used off-label to treat chronic pain and various systemic inflammatory and autoimmune conditions (Bostick, McCarter, & Nykamp, 2019; Carvalho & Skare, 2023; Ekelem, Juhasz, Khera, & Mesinkovska, 2019; Li, You, Griffin, Feng, & Shan, 2018; Miskoff & Chaudhri, 2018; Qu, Meng, Handley, Wang, & Shan, 2021; Toljan & Vrooman, 2018). In addition to its pain modulation, early evidence has suggested that low-dose NTX may be a promising anticancer treatment option (Li et al., 2018; Liubchenko et al., 2021). One proposed mechanism of action for its efficacy in cancer therapy involves the blockage of the opioid growth factor ([Met^5^]-enkephalin, OGF) and its receptor (OGFr), a regulatory axis suggested to inhibit cell proliferation in human cancer and normal cells (Donahue, McLaughlin, & Zagon, 2009; Ian S. Zagon, Donahue, & McLaughlin, 2009), through modulation of the G1/S cell cycle phase via cyclin-dependent kinase inhibitory pathways (F. Cheng, McLaughlin, Verderame, & Zagon, 2008, 2009; F. Cheng, Zagon, Verderame, & McLaughlin, 2007). Notably, OGFr is suggested to be markedly reduced in human head and neck squamous cell carcinoma, with tumor tissue exhibiting ninefold fewer OGFr binding sites and a fivefold reduction in OGFr protein levels compared to controls (Patricia J. McLaughlin, Stack, Levin, Fedok, & Zagon, 2003), implicating a dysregulation of this modulatory pathway in cancer disease progression (Fanning et al., 2012; Ian S. Zagon et al., 2009). While in-vitro and animal studies have suggested that NTX may restore the functionality of OGF-OGFr axis and suppress tumor growth (F. Cheng et al., 2009; Donahue, McLaughlin, & Zagon, 2011c; Ian S. Zagon, Verderame, & McLaughlin, 2003), findings remain controversial regarding dosing, treatment schedule, and mechanism of action (Liubchenko et al., 2021).

To address these gaps, the primary aim of this study was to investigate the efficacy of various doses, durations, and schedules of NTX on the viability of OSCC in comparison to control cells and variations of OGF-OFGr expression as a function of tumor stage and NTX treatment. Based on previous literature (Donahue, McLaughlin, & Zagon, 2011b; Patricia J. McLaughlin, Levin, & Zagon, 2003), we hypothesized that treatment of OSCC with intermittent administration of NTX would inhibit OSCC cell growth. We also hypothesized that OGF-OGFr expression would differ between OSCC and control cells and would be modulated as a result of NTX therapy. We also hypothesized that NTX may sensitize OSCC cells to other therapies. Thus, a secondary aim was to analyze the efficacy of NTX in combination with standard cytotoxic chemotherapeutic agents and its effect on short-term changes in OGFr expression in treated cells. Finally, to test for differences in NTX response between tumor stages, we chose two cell line models representing locally invasive and metastatic carcinoma: SCC-25 and Detroit 562, respectively. We have previously shown that these lines differ in their biomarker expression and drug response, supporting their use in the present study (Hamoui, Rizvi, Arnouk, & Roberts, 2025).

## 2. Materials and Methods

### 2.1. Chemicals, drugs, and reagents

Naltrexone (NTX) and Sulphorhodamine B (SRB) were purchased from MilliporeSigma (Burlington, MA, USA). Cisplatin was purchased from Avantor (Radnor, PA, USA). Docetaxel and Cell Counting Kit 8 (CCK-8) reagents were acquired from Selleck Chemicals (Houston, TX, USA). Additional CCK-8 reagent was purchased from GlpBio (Montclair, CA, USA). Hydrocortisone for growth media supplementation was purchased from STEMCELL Technologies (Cambridge, MA, USA).

### 2.2. Cell lines and cell culture

In order to study two different stages of OSCC, two oropharyngeal carcinoma lines were selected. SCC-25 was derived from a locally invasive squamous cell carcinoma on the tongue of a 70-year-old male patient (Hamoui et al., 2025). These cells were grown in DMEM-F12 media supplemented with 400 ng/mL hydrocortisone, 10% fetal bovine serum (FBS), and 1% penicillin-streptomycin (P/S). Detroit-562 was derived from a pleural effusion of a female patient with metastatic pharyngeal carcinoma, and therefore represents a more advanced disease (Hamoui et al., 2025). These cells were maintained in EMEM supplemented with 10% FBS and 1% P/S. Normal primary gingival keratinocytes (PGK) were grown in Dermal basal media supplemented with a Keratinocyte Growth Kit from the American Type Culture Collection (ATCC, Manassas, VA, USA) and 1% P/S. SKOV-3 ovarian carcinoma cells were selected as positive controls for the effects of naltrexone on the basis of prior studies (Donahue, McLaughlin, & Zagon, 2011a; Donahue et al., 2011b; Ian S. Zagon et al., 2009). SKOV-3 were grown in RPMI plus 10% FBS and 1% P/S. SKOV-3 cells were acquired from ATCC. PGK, SCC-25, and Detroit 562 cells were provided by Dr. Hilal Arnouk at Midwestern University. Cell cultures were maintained in a 37°C humidified incubator with 5% CO2 atmosphere and used within 12 passages of thaw to prevent drift.

### 2.3. Sulphorhodamine B assays

Effects of NTX as a single agent were evaluated by SRB assay. Briefly, cells were plated at 1,500-3,000 cells per well in 96-well plates. The following day, media was removed and replaced by NTX at 1 µM or 10 µM in normal media, or plain media as a control. NTX was added and removed as outlined in **Table 1**, yielding treatments designated 5h once, 5h daily, 5h every other day (EOD), and constant exposure. After 72 hours, media and any remaining NTX were removed and cells were fixed in 100 µl per well 10% trichloroacetic acid (TCA) at 4°C for one hour. TCA was then removed and cells were washed with 200 µl water and allowed to air dry. Fixed cells were then stained with 0.4% SRB in 1% acetic acid for 15 min at room temperature. SRB stain was discarded and wells were rinsed 3-4 times with 1% acetic acid until no further pink color was present in the discarded wash. Stray SRB around the walls of the wells was removed with a fine-point cotton swab, taking care not to disturb stained cells on the bottom of the wells. SRB was solubilized in 200 µl per well 10 mM Tris base, pH 10.5, and plates were read for absorbance at 570 nm in an EnSpire Plate reader (Perkin Elmer, Waltham, MA, USA) to quantify SRB signal. All experiments were repeated at least three times with 5 technical replicates in each run.

**Table 1.** Treatment schemes for NTX single agent experiments.

|  | Day 1 | Day 2 |  | Day 3 |  | Day 4 |  | Day 5 |
| --- | --- | --- | --- | --- | --- | --- | --- | --- |
|  |  | NTX<br>On | NTX<br>Off | NTX<br>On | NTX<br>Off | NTX<br>On | NTX<br>Off |  |
| **Control** | Plate cells | - | - | - | - | - | - | SRB Assay |
| **Constant** |  | + | - | + | - | + | - |  |
| **5h Once** |  | + | + | - | - | - | - |  |
| **5h Daily** |  | + | + | + | + | + | + |  |
| **5h EOD** |  | + | + | - | - | + | + |  |

NTX: naltrexone; SRB: Sulphorhodamine B.
Constant: 72 hours of continuous exposure;
5h Once: one 5h-exposure;
5h Daily: 5h treatment daily for three days;
EOD: 5h treatment every other day.

### 2.4. Cell Counting Kit 8 assays

Efficacy of combination therapy was evaluated via Cell Counting Kit 8 (CCK-8) assay. Briefly, cells were plated at 3,000 cells per well in 96-well plates. The following day, cells were treated with 10 µM NTX following the dosing regimens in **Table 2**, or plain media control in 100 µl total volume. Wells with media only and no cells were used for background subtraction. The following day, cisplatin or docetaxel was added in an additional 100 µl, bringing total well volume to 200 µl. Cells were incubated for 48 hours, and then 20 µl per well of CCK-8 reagent was added, including to background wells. After a further 2 hour incubation, plates were read at 450 nm in a BioTek Synergy H1 plate reader (Agilent Technologies, Santa Clara, CA, USA). Raw data were analyzed using Microsoft Excel, subtracting the average reading from the background wells from all sample values and normalizing to untreated control wells. All experiments were repeated at least three times with 3 technical replicates per run.

**Table 2.** Treatment schemes for drug combination experiments.

|  | Day 1 | Day 2 |  | Day 3 |  | Day 4 | Day 5 |
| --- | --- | --- | --- | --- | --- | --- | --- |
|  |  | NTX<br>On | NTX<br>Off | NTX<br>Off | Chemo<br>On |  |  |
| **Control** | Plate cells | - | - | - | + | Incubate | CCK-8 Assay |
| **Constant** |  | + | - | - | + |  |  |
| **5h** |  | + | + | - | + |  |  |
| **24h** |  | + | - | + | + |  |  |

CCK-8: Cell Counting Kit 8; Chemo: chemotherapeutic agents (cisplatin or docetaxel); NTX: naltrexone.
5h: 5-hour pretreatment followed by 19 hours normal media;
24h: 24-hour pretreatment;
Constant: continuous treatment (24 hour pretreatment + continued NTX treatment during chemotherapy exposure).

### 2.5. Western Blotting

Cells with and without NTX treatment were harvested using trypsin and pelleted. Cell pellets were lysed in RIPA buffer supplemented with protease and phosphatase inhibitors and PMSF. Protein in lysates was quantified by BCA assay (ThermoFisher Scientific, Waltham, MA, USA) and equal masses and volumes of protein were run on TGX precast gradient gels from Bio-Rad (Hercules, CA, USA). Protein was then transferred to PVDF membranes using the Transblot SD semi-dry transfer system from Bio-Rad. Membranes were blocked in 5% milk in phosphate buffered saline with 0.5% Tween 20 (PBST) for 40 min at room temperature before incubation with primary antibodies in 5% bovine serum albumin in PBST overnight at 4°C. Blots were then washed in PBST, incubated with secondary antibodies in 5% milk in PBST for one hour at room temperature, washed again, and imaged using Clarity ECL substrate and a ChemiDoc imager, both from Bio-Rad. Band intensities were quantified using Bio-Rad Image Lab software, version 5.2.1. Primary antibodies were anti-OGFr (1:1,000, #11177-1-AP from Proteintech, Rosemont, IL, USA) and anti-beta actin (1:500, #sc-47778 from Santa Cruz Biotechnology, Dallas, TX, USA). Secondary antibodies for OGFr and Actin, respectively, were horse anti-rabbit (1:5,000, #7074) and anti mouse (1:10,000, #7076) from Cell Signaling Technology (Danvers, MA, USA).

## 3. Results

### 3.1. OSCC cells express OGFR

At first, we verified that OGFr was expressed in our OSCC lines. We did this by comparing them to normal human Primary Gingival Keratinocytes (PGK) cells, and saw that SCC-25 had similar OGFr protein expression, while Detroit 562 cells had somewhat higher levels of OGFr (n=1, **Supplementary Figure S1a**). This could be because the Detroit 562 cell line is a pharyngeal carcinoma line derived from metastatic tissue, which has increased proliferation and altered regulatory pathways causing dysregulation.

### 3.2. Low dose naltrexone does not significantly impact OSCC cell survival

To evaluate the efficacy of NTX, two different doses were tested in two distinct head and neck cancer cell lines: SCC-25 and Detroit 562. Each NTX dose was used in a scheduling of constant 72 hour exposure (Const), single 5h treatment (Once), 5h treatment daily for three days (Daily), or 5h treatment every other day (EOD). Any inhibition of growth was not repeatable, and no statistically significant changes in cell survival were observed in either SCC-25 (n=4, **Figure 1a**) or Detroit 562 (n=3, **Figure 1b**).

**Figure 1.**
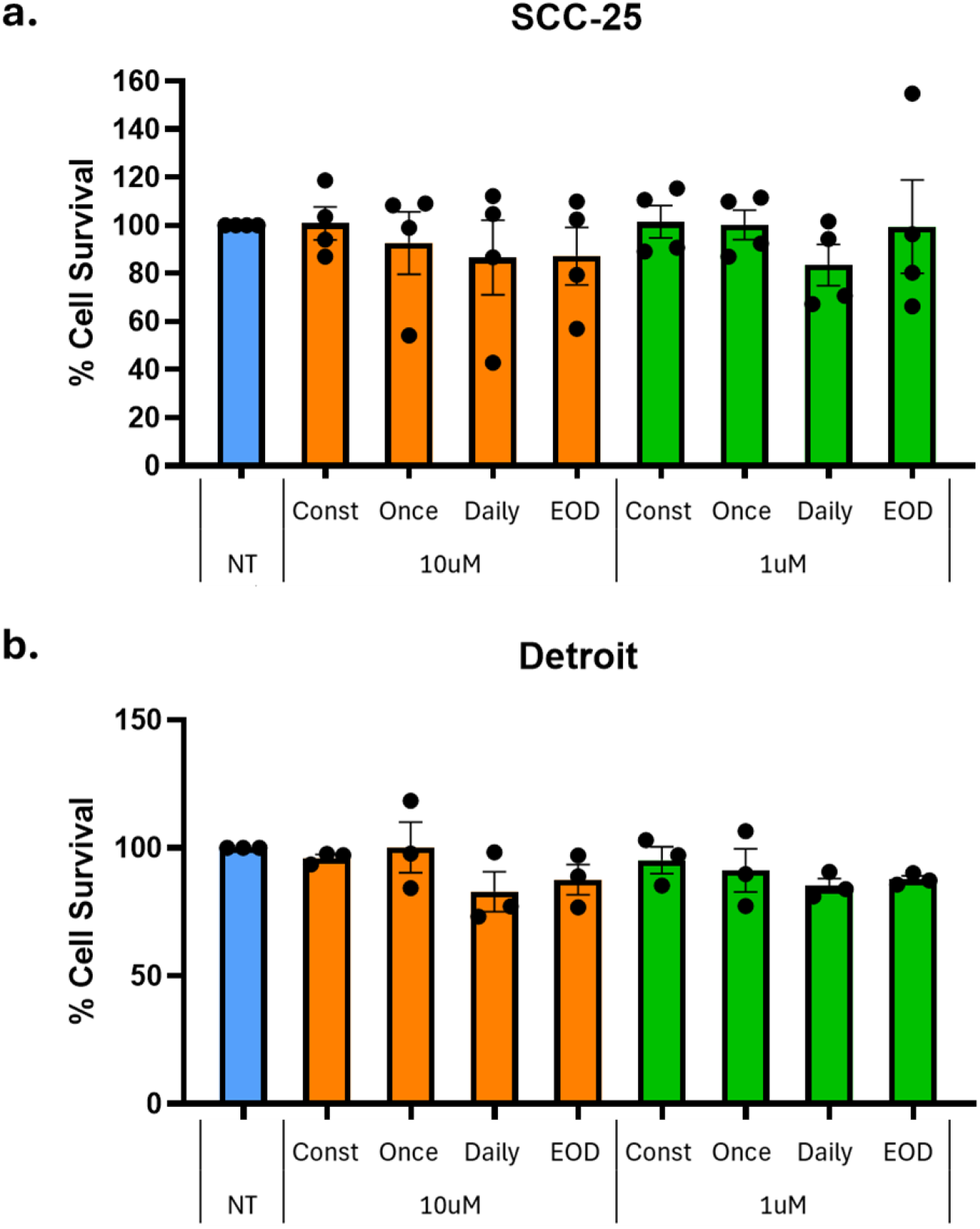
Evaluation of NTX efficacy in cell lines. Two doses were tested in **a.** SCC-25 and **b**. Detroit 562 cells. Each dose was used with constant 72 hour exposure (Const), single 5h treatment (Once), 5h treatment daily for three days (Daily), or 5h treatment every other day (EOD). NT, no treatment control. Any inhibition of growth was not repeatable, and no statistically significant changes in cell survival were observed (one-way ANOVA, SCC-25 n=4, Detroit 562 n=3, bars represent mean ± SEM). Dots represent individual values.

### 3.3. Naltrexone has mild effects on SKOV-3 ovarian cancer cells

With previous literature proving that NTX affects growth of SKOV-3 ovarian cancer cells (Donahue et al., 2011c), we repeated our experiments in this cell line to assess the efficacy of our dosage and timing schedules of NTX treatment. We first verified that OGFr expression was similar in this line (n=1, **Supplementary Figure S1b**). When SKOV-3 was plated at low density, using equal starting cell number to OSCC cells, constant NTX drove cell growth, while intermittent dosing (i.e., once, daily, or EOD) had a slight suppressive effect, up to 10µM, though these changes were not statistically significant (**Figure 2a**). At higher doses, all dosing schemes were toxic (n=1, **Supplementary Figure S2**). However, we noted SKOV-3 cells grew more slowly than the OSCC lines, which may have affected our ability to detect significant changes. We therefore repeated the experiment with 3,000 cells plated per well, giving ending density similar to the OSCC lines. Variability increased, and no statistically significant changes were observed, though some replicates led to reduced growth in the presence of intermittent NTX, particularly EOD (**Figure 2b**). While not completely transparent results, these findings do suggest that the efficacy and cellular response to NTX in SKOV-3 cells and OSCC cells are similar, and may be influenced by both dosing strategy and cell density, highlighting the importance of validating methods when evaluating NTX treatment outcomes.

**Figure 2.**
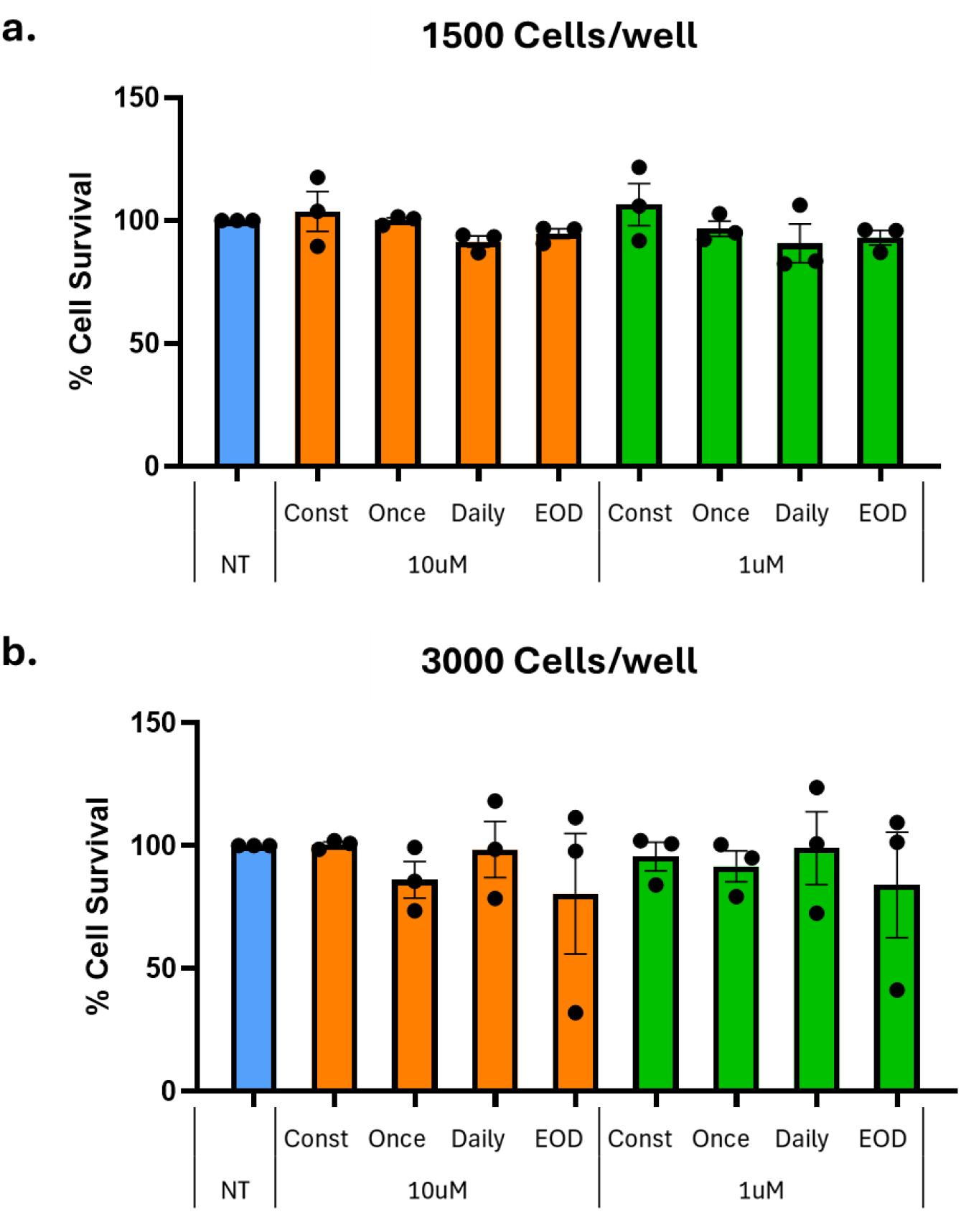
Effects of NTX dosing on SKOV-3 ovarian cancer cells. **a.**At low density, effect size is low, but there is a slight trend toward growth with constant exposure, and suppression with intermittent dosing. **b**. Higher cell density leads to greater variability, but some replicates do show marked reduction in SKOV-3 growth with intermittent NTX. n=3 for both densities and graphs show mean ± SEM. Dots represent individual values.

### 3.4. Naltrexone enhances the efficacy of chemotherapy in OSCC cells

To evaluate the potential of naltrexone (NTX) as an adjunct to standard chemotherapy, SCC-25 and Detroit 562 cells were treated with 10 µM NTX under three exposure schedules: 5-hour pretreatment followed by 19 hours normal media, 24-hour pretreatment, and continuous treatment (24 hour pretreatment plus continued NTX presence during chemotherapy exposure). After the initial 24 hour pretreatment window, cells were treated with increasing concentrations of either cisplatin or docetaxel, and viability was assessed after 48 hours. Across both cell lines, NTX treatment resulted in decreased cell survival compared to chemotherapy alone, with short-term exposures (5-hour and 24-hour) generally producing stronger effects than continuous treatment (**Figure 3**). The degree of sensitization varied depending on both the chemotherapy agent used and the NTX exposure. NTX sensitized SCC-25 cells to cisplatin, particularly when 24 h pretreatment was used (**Figure 3a**). In Detroit 562 cells, significant changes from no treatment controls were only observed with low or absent levels of cisplatin (**Figure 3b**). In both cell lines, treatment with 5h NTX prior to docetaxel was more efficacious than docetaxel alone (**Figure 3c**,**d**). These results suggest that NTX may increase the cytotoxic effect of chemotherapeutic agents in a cell line dependent and specific manner. Shorter NTX exposures appeared to be more effective than continuous treatment, indicating that timing plays a critical role in maximizing the benefit. This supports the potential utility of NTX as a chemosensitizing agent in OSCC therapy.

**Figure 3.**
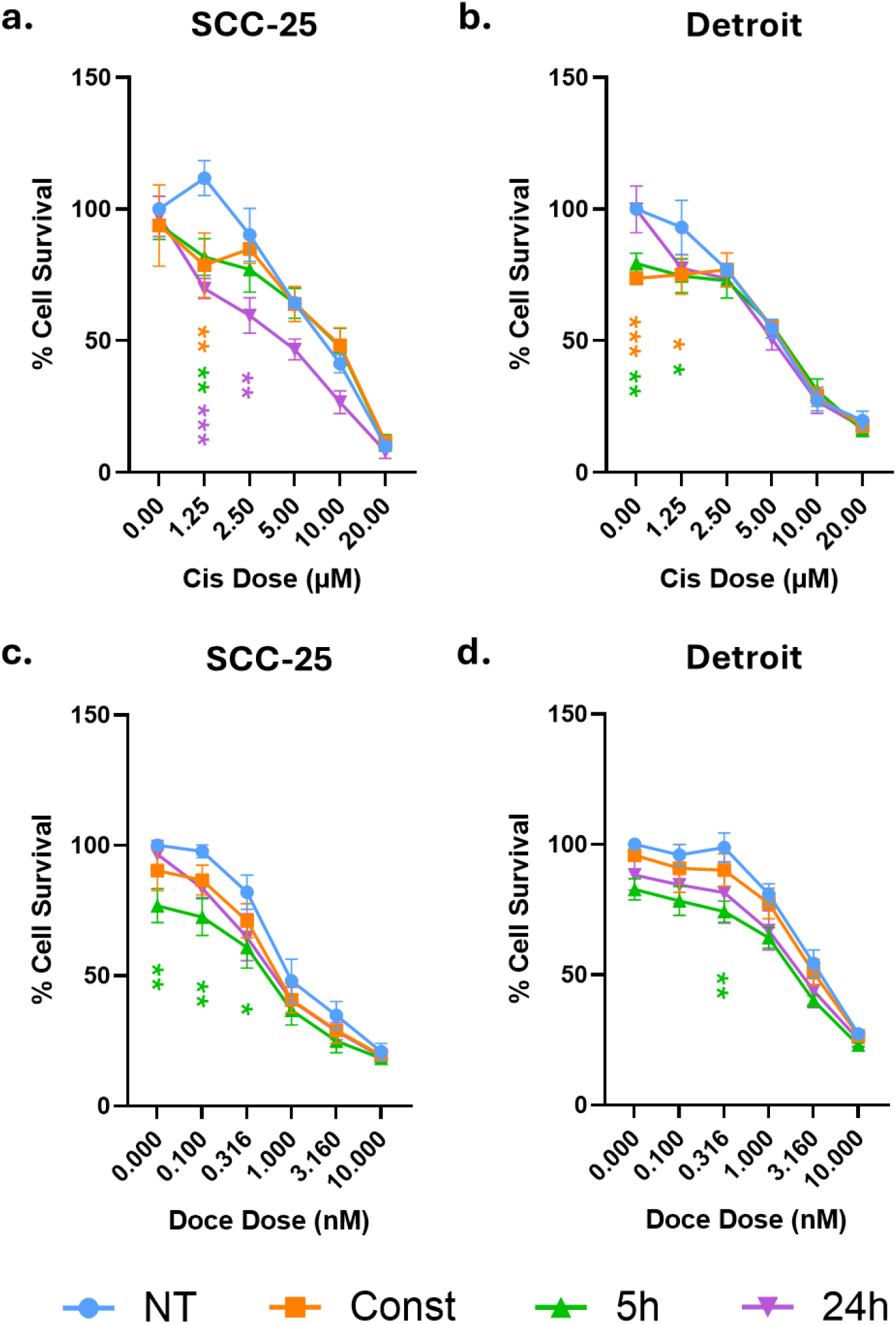
Effects of combining NTX with chemotherapy.**a.**In SCC-25, NTX sensitized cells to cisplatin (Cis), with 24h exposure performing the best. **b**. Combination of NTX with Cis in Detroit 562 cells. Constant and 5h exposure to NTX had a slight sensitizing effect. **c**,**d**. Combination of NTX with docetaxel (Doce) was more effective than Doce alone in both SCC-25 (**c**.) and Detroit 562 (**d**.) cells. 5h exposure had a statistically significant effect in both lines. Data were analyzed with two-way ANOVA comparing conditions at each Cis/Doce dose. Graphs show mean ± SEM of n=3 (Cis) or n=4 (Doce) independent experiments. ^*^ p<.05, ^**^ p<.01, ^***^ p<.001. Asterisks represent condition of same color versus NT at that dose.

### 3.5. Effect of naltrexone exposure on OGFr expression is cell line dependent

OGFr expression was compared in untreated OSCC cells, those treated with 10µM NTX once for 5 h followed by 19 h of normal media, and those treated with constant NTX exposure for the full 24 h. In SCC-25 cells, there was a trend toward increased OGFr expression with NTX treatment duration, but due to inter-experiment variation, the difference did not achieve statistical significance (**Figure 4a**,**b**, n=4). No clear trend was observed for OGFr expression following NTX exposure in Detroit 562 cells (**Figure 4a**,**c**, n=4).

**Figure 4.**
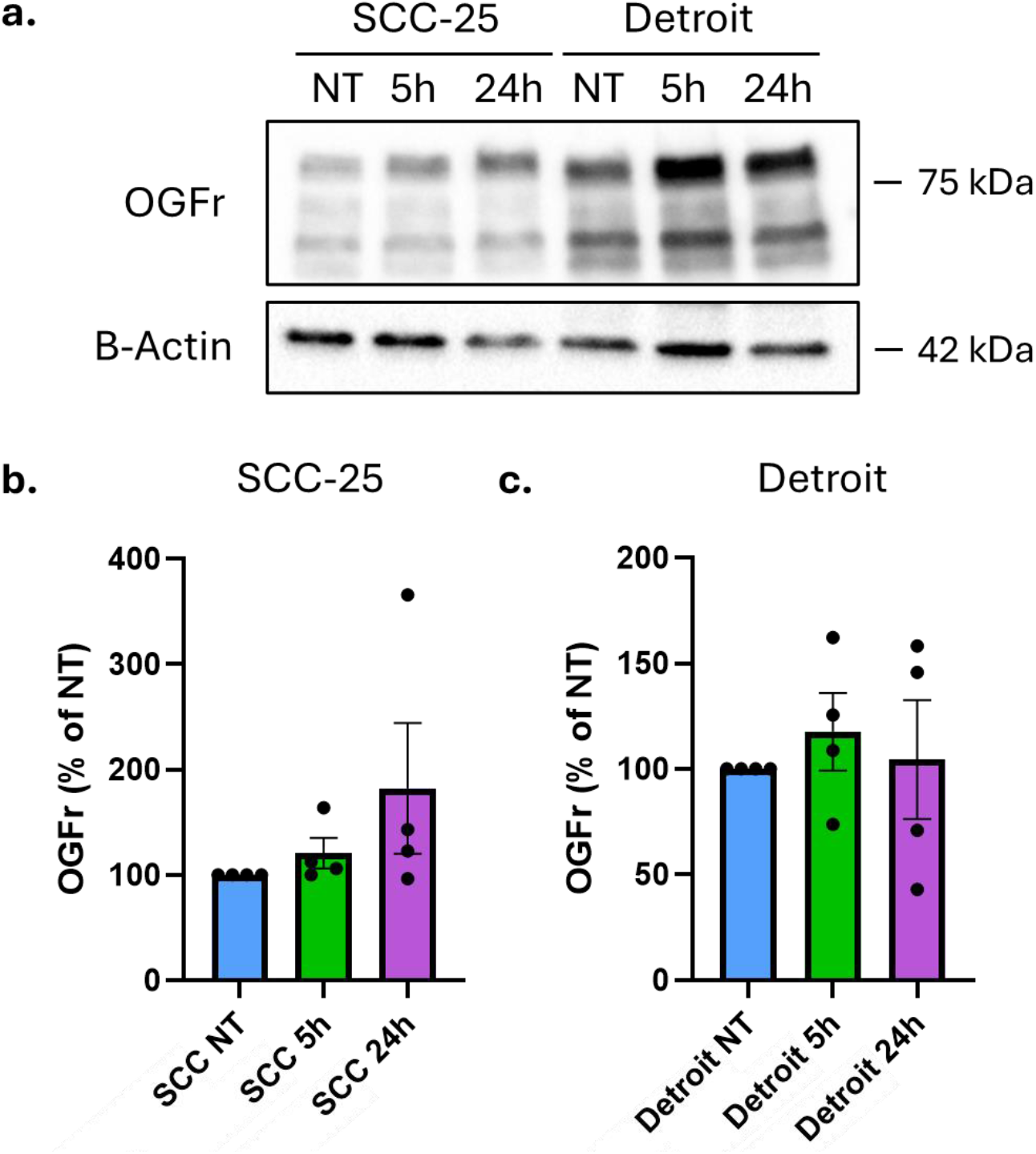
Evaluation of OGFr expression in cell lines. **a.**OGFr expression was compared in untreated SCC-25 cells (NT), those treated with 10µM NTX for five hours (5h), and those treated for 24 hours (24h). Representative blot shown. **b**. Quantification of n=4 replicate experiments shows a trend toward increased OGFr expression in NTX-treated SCC-25 cells that did not achieve statistical significance due to high variability. **c**. Detroit 562 cells treated with NTX do not show a consistent trend in OGFr expression changes (n=4). Graphs show mean ± SEM. Dots represent individual values.

## 4. Discussion

OSCC remains a major clinical challenge, particularly when diagnosed at an advanced stage, when treatment options are limited and outcomes are poor. There is growing need for adjuvant therapies that can be proposed as standalone management options or potentiate existing treatments without introducing substantial toxicity. In this study, we evaluated the effect of NTX, alone and in combination with existing chemotherapeutic agents, on OSCC cell viability and OFGr expression. We selected two cell lines representing different stages of OSCC disease progression and evaluated responses under different NTX exposure regimens.

### 4.1. Minimal effect of NTX on OSCC cell survival

As a monotherapy, NTX demonstrated minimal impact on OSCC cell viability, with no consistent or reproducible inhibition of cell proliferation observed across experiment replicates or cell lines (SCC-25 or Detroit 562), regardless of the NTX exposure regimen (**Figure 1**). A recent systematic review (Liubchenko et al., 2021) highlighted the heterogeneity of NTX exposure regimens across in vitro cancer cell studies, where NTX has been administered intermittently, continuously, or for short-term duration from 72 to 120 hours, at concentrations between 10^−6^ and 10^−5^. In the current study, we implemented several exposure exposure schedules - continuous treatment for 72 hours, a single 5-hour exposure, repeated daily 5-hour exposure over three days, or every-other-day 5-hour administration. These treatment schedules were informed by previous studies suggesting that intermittent or short-term NTX may be more effective than continuous treatment in inhibiting tumor growth compared to controls (Liubchenko et al., 2021; Patricia J. McLaughlin & Zagon, 2015). Indeed, previous studies have shown that short-term NTX exposure can inhibit tumor cell growth by 24% to 42% across multiple cancer cell lines (Liubchenko et al., 2021), including SKOV-3 and OVCAR-3 (ovarian cancer), MDA-MD-231 (triple-negative breast cancer), SCC-1 (oral squamous carcinoma), HCT-116 (colon carcinoma), and MIA PaCa-2 (pancreatic carcinoma) (Donahue et al., 2011b, 2011a; Ian S Zagon, Porterfield, & McLaughlin, 2012). Conversely, continuous NTX treatment has been associated with increased proliferation (ranging from 9 to 71%) in various human and murine cancer types, possibly due to sustained opioid receptor blockade (Donahue et al., 2009, 2011b; Donahue, McLaughlin, & Zagon, 2011d; P J McLaughlin, Levin, & Zagon, 1999; Patricia J McLaughlin et al., 2009; I. S. Zagon, Hytrek, & McLaughlin, 1996; Ian S. Zagon, 1988; I.S. Zagon & McLaughlin, 1990).

To the best of our knowledge, only two studies so far have directly investigated the effect of NTX in OSCC, both from the same research group. These studies have observed a 26% reduction in tumor cell proliferation compared to control with short-term NTX (72 hours), while continuous treatment over the same duration resulted in a comparable increase in proliferation (Donahue et al., 2011b; Ian S. Zagon et al., 2009). Possible explanations for this discrepancy in results may be attributed to the different cell lines utilized in the studies. Although all classified as OSCC, these cell lines are biologically and genetically different. For example, SCC-25 (used in our study) and SCC-1 (used by Donahue et al. and Zagon et al.) are both tongue squamous carcinoma cell lines, but differ in p53 status, OGFr expression, and OGF-OGFr axis activity, which could influence response to NTX (Dudás et al., 2018; Sano et al., 2011). Similarly, Detroit 562 originates from pharyngeal carcinoma and manifests as more aggressive and less differentiated. Cell culture conditions, media supplements, drug stability and dosing precision could have also influenced the different response to NTX treatment.

### 4.2. NTX effect may vary according to cell line

Given consistent results across multiple studies, we applied NTX treatment to SKOV-3 ovarian cancer cells. When SKOV-3 cells were plated at low density, continuous NTX exposure appeared to enhance cell proliferation (although mild and only up to 10 µM in some experiments), while an intermittent dosing schedule slightly suppressed cellular growth. These findings are consistent with existing studies in the literature (Donahue et al., 2011a, 2011b; Stockdale et al., 2017). Conversely, higher NTX concentrations were toxic regardless of the dosing regimen, aligning with known off-target toxicity at elevated NTX concentrations (France et al., 2021). In contrast, when these experiments were repeated using higher cell density (thus comparable to OSCC cell confluency), an opposite trend was observed, characterized by reduced or absent stimulatory effect with continuous dosing schedule but greater growth suppression with intermittent dosing. These results suggest that not only cell proliferation may be influenced by NTX dosing schedule, but cell density may also modulate SKOV-3 response to NTX, possibly due to density-related changes in paracrine signaling, receptor availability, or metabolic state. Moreover, previous studies have shown that the growth-modulating effect of NTX in SKOV-3 is fully suppressed when OGFr is silenced via siRNA, suggesting that the OGF-OGFr axis is the sole mediator of NTX action in this cell line (Donahue et al., 2009). This finding may explain a higher observed reproducibility of cell behavior with SKOV-3 compared to OSCC cell lines, where alternative or competing pathways may influence NTX response. Future studies may focus on further analyzing these dependencies and optimizing our regimens to yield efficacy more in line with previous reports.

### 4.3. Enhanced effect of NTX in combination with chemotherapeutic agents

Chemotherapeutic agents, such as cisplatin and docetaxel, are commonly used in the treatment of OSCC, especially for locally advanced disease or in advanced stages with regional or widespread metastasis. Nevertheless, their use is limited by numerous side effects, including nephrotoxicity, peripheral neuropathy, nausea, vomiting, and ototoxicity (Elmorsy et al., 2024). Such side effects are dose-dependent; thus, exploration of combination therapies that allow to reduce the dosage - and therefore the toxicity - of chemotherapeutic agents is encouraged (Dasari & Tchounwou, 2014). In our study, the consistent leftward and downward shift of the cell viability curves under NTX conditions relative to controls without NTX treatment suggests that NTX potentiate the cytotoxic efficacy of both cisplatin and docetaxel in SCC-25 and Detroit 562 cells. Short-term NTX exposure (5-hour and 24-hour) generally resulted in lower cell viability compared to chemotherapy alone or continuous NTX regimen, with the exception of Detroit 562 treated with low-dose cisplatin (0-1.25 µM). Interestingly, at higher concentrations of cisplatin (5-20 µM), cisplatin alone produced greater cytotoxicity than when combined with either 5-hour or continuous NTX exposure. Our findings are consistent with similar in-vitro studies conducted in lung, colorectal carcinoma, and ovarian cancer, where NTX regimen - such as single 48-hour or repeated 6-hour NTX treatment every other day over 5 days - resulted in a 20-45% tumor growth reduction in viability compared to chemotherapy alone (i.e., cyclophosphamide, gemcitabine, taxol, cisplatin, and oxaliplatin) (Donahue et al., 2011a; Liu et al., 2016). Overall, these findings support a dose-enhancing effect of NTX on chemotherapeutic cytotoxicity, particularly under intermittent exposure conditions, which appeared to be the most effective regimen. This dose-enhancing effect may be mediated through the interaction between NTX and OGF-OGFr axis on modulation of cell proliferation. By modulating this pathway, NTX may sensitize cells to the DNA-damaging effects of chemotherapeutic agents, resulting in a synergistic cytotoxicity, cell apoptosis, suppression of cell cycle, and promoting an immunogenic microenvironment at lower chemotherapeutic doses (Avella et al., 2010; Donahue et al., 2011a; Ma, Wang, Liu, Shan, & Feng, 2020). These results may contribute to the body of combination therapies that enhance tumor cell growth inhibition while allowing for lower doses of chemotherapeutic agents (Mokhtari et al., 2017).

### 4.4. OGFr expression and modulation by NTX

In addition to its known binding capacity to μ-, δ- and κ-opioid, NTX also targets the OGFr by competitively inhibiting the binding of OGF. The OGF-OGFr complex modulates cell proliferation by upregulating cyclin-dependent kinase inhibitors p16 and p21, resulting in decreased phosphorylation of retinoblastoma protein (Rb) and subsequent delay of the G1 to S phase transition in the cycle phase (F. Cheng et al., 2009, 2007; Donahue et al., 2011c). NTX competes with OGF for binding to OGFr, thus disrupting OGF-OGFr interaction. Notably, this disruption has paradoxical effects that are highly dependent on NTX dosing schedule, thereby intermittent or low-dose NTX has been shown to enhance OGF-OGFr signaling via a compensatory rebound mechanism and inhibit tumor growth, whereas continuous NTX exposure may suppress OGFr expression and promotes cell growth (Liubchenko et al., 2021). Our results suggested that OFGr is expressed in both SCC-25 and Detroit 562 cell lines, with slightly higher baseline expression in Detroit 562 compared to normal controls. Various research has examined the OGF-OGFr axis across different cancer types, with some studies supporting a dysregulation of the pathway, especially in more aggressive and less differentiated tumors, and others reporting that the axis is maintained intact or only minimally altered (Avella et al., 2010; Patricia J. McLaughlin, Stack, et al., 2003; Patricia J McLaughlin & Zagon, 2006). Thus, alterations of the OGF-OGFr axis appear to be tumor-type specific and potentially influenced by cancer stages (Patricia J McLaughlin & Zagon, 2006; Ian S Zagon & McLaughlin, 2006). Interestingly, our results showed that continuous NTX exposure resulted in a modest increase in OFGr expression in SCC-25 cells, with inconsistent effects in Detroit 562. This contrasts with the idea that sustained opioid receptor blockade can downregulate OGFr (Donahue et al., 2011b). Advanced stage cells (i.e., Detroit 562) showing higher basal levels of OGFr than earlier stage (SCC-25) differs from prior reports and may partially explain our findings (Patricia J McLaughlin & Zagon, 2006). Moreover, in contrast to previous studies showing that intermittent NTX exposure enhances OGF-OGFr signaling and suppressing proliferation, we did not observe a consistent upregulation of OGFr expression under those conditions, highlighting the need for further investigation.

### 4.5. Limitations

The current study has several limitations. As all experiments were done in cell lines, not all factors present in human patients are accounted for. For example, no paracrine or microenvironment effects on tumor growth were accounted for in our models. In addition, statistical significance was impeded by the variability of our results, and further study will be needed to identify the source(s) of this variation. Furthermore, due to the number of dosing schemes and drug combinations tested, uniform total treatment duration was used for all experiments. In future, NTX treatments between 5 and 24 hours may be tried, with overall assay durations greater than the 48-72 hours used here. Finally, additional studies will be necessary to determine the mechanism of NTX action on OSCC cells. OGFr levels were not significantly changed, especially in Detroit 562 cells, but the activity of the receptor and its downstream signals may still be impacted, as outlined above.

## Author Contributions

L.S., F.G., C.M.R. conceptualized the study; F.G. and C.M.R. developed the study methodology; E.S., S.K., C.M.R., Z.A.R. conducted the investigation; C.M.R. performed formal analysis; L.S., E.S., S.K., C.M.R. acquired funding for the study; C.M.R. supervised the project; L.S., F.G., C.M.R., E.S., and S.K. prepared the original draft of the manuscript; all authors reviewed and edited the manuscript. All authors approved the current version of the manuscript.

## Acknowledgements

We thank Dr. Hilal Arnouk who provided PGK, SCC-25, and Detroit 562 cells. We gratefully acknowledge Ellen Kohlmeir and the MWU Core Facility. BCA, CCK-8, and SRB assays were performed using the PerkinElmer EnSpire Plate Reader, and cells were counted using the Denovix CellDrop automated cell counter located in the Core. We also gratefully acknowledge the financial support of the MWU College of Dental Medicine. This project was also financially supported by the MWU College of Dental Medicine Dean’s Research Fellowship to ES and laboratory startup funding to CMR.

## References

Avella, D. M., Kimchi, E. T., Donahue, R. N., Tagaram, H. R. S., McLaughlin, P. J., Zagon, I. S., & Staveley-O’Carroll, K. F. (2010). The opioid growth factor-opioid growth factor receptor axis regulates cell proliferation of human hepatocellular cancer. American Journal of Physiology-Regulatory, Integrative and Comparative Physiology, 298(2), R459–R466. 10.1152/ajpregu.00646.2009

Barsouk, A., Aluru, J. S., Rawla, P., Saginala, K., & Barsouk, A. (2023). Epidemiology, Risk Factors, and Prevention of Head and Neck Squamous Cell Carcinoma. Medical Sciences, 11(2), 42. 10.3390/medsci11020042

Beckham, T. H., Leeman, J. E., Xie, P., Li, X., Goldman, D. A., Zhang, Z., … Tsai, C. J. (2019). Long-term survival in patients with metastatic head and neck squamous cell carcinoma treated with metastasis-directed therapy. British Journal of Cancer, 121(11), 897–903. 10.1038/s41416-019-0601-8

Bostick, K. M., McCarter, A. G., & Nykamp, D. (2019). The Use of Low-Dose Naltrexone for Chronic Pain. The Senior Care Pharmacist, 34(1), 43–46. 10.4140/tcp.n.2019.43

Chamoli, A., Gosavi, A. S., Shirwadkar, U. P., Wangdale, K. V., Behera, S. K., Kurrey, N. K., … Mandoli, A. (2021). Overview of oral cavity squamous cell carcinoma: Risk factors, mechanisms, and diagnostics. Oral Oncology, 121, 105451. 10.1016/j.oraloncology.2021.105451

Cheng, F., McLaughlin, P. J., Verderame, M. F., & Zagon, I. S. (2008). The OGF-OGFr axis utilizes the p21 pathway to restrict progression of human pancreatic cancer. Molecular Cancer, 7(1), 5. 10.1186/1476-4598-7-5

Cheng, F., McLaughlin, P. J., Verderame, M. F., & Zagon, I. S. (2009). The OGF–OGFr Axis Utilizes the p16INK4a and p21WAF1/CIP1 Pathways to Restrict Normal Cell Proliferation. Molecular Biology of the Cell, 20(1), 319–327. 10.1091/mbc.e08-07-0681

Cheng, F., Zagon, I. S., Verderame, M. F., & McLaughlin, P. J. (2007). The Opioid Growth Factor (OGF)–OGF Receptor Axis Uses the p16 Pathway to Inhibit Head and Neck Cancer. Cancer Research, 67(21), 10511–10518. 10.1158/0008-5472.can-07-1922

Cheng, Y., Li, S., Gao, L., Zhi, K., & Ren, W. (2021). The Molecular Basis and Therapeutic Aspects of Cisplatin Resistance in Oral Squamous Cell Carcinoma. Frontiers in Oncology, 11, 761379. 10.3389/fonc.2021.761379

Dasari, S., & Tchounwou, P. B. (2014). Cisplatin in cancer therapy: Molecular mechanisms of action. European Journal of Pharmacology, 740, 364–378. 10.1016/j.ejphar.2014.07.025

de Carvalho, J. F., & Skare, T. (2023). Low-Dose Naltrexone in Rheumatological Diseases. Mediterranean Journal of Rheumatology, 34(1), 1. 10.31138/mjr.34.1.1

Donahue, R. N., McLaughlin, P. J., & Zagon, I. S. (2009). Cell proliferation of human ovarian cancer is regulated by the opioid growth factor-opioid growth factor receptor axis. American Journal of Physiology-Regulatory, Integrative and Comparative Physiology, 296(6), R1716– R1725. 10.1152/ajpregu.00075.2009

Donahue, R. N., McLaughlin, P. J., & Zagon, I. S. (2011a). Low-dose naltrexone suppresses ovarian cancer and exhibits enhanced inhibition in combination with cisplatin. Experimental Biology and Medicine, 236(7), 883–895. 10.1258/ebm.2011.011096

Donahue, R. N., McLaughlin, P. J., & Zagon, I. S. (2011b). Low-dose naltrexone targets the opioid growth factor–opioid growth factor receptor pathway to inhibit cell proliferation: mechanistic evidence from a tissue culture model. Experimental Biology and Medicine, 236(9), 1036–1050. 10.1258/ebm.2011.011121

Donahue, R. N., McLaughlin, P. J., & Zagon, I. S. (2011c). The opioid growth factor (OGF) and low dose naltrexone (LDN) suppress human ovarian cancer progression in mice. Gynecologic Oncology, 122(2), 382–388. 10.1016/j.ygyno.2011.04.009

Donahue, R. N., McLaughlin, P. J., & Zagon, I. S. (2011d). Under-expression of the opioid growth factor receptor promotes progression of human ovarian cancer. Experimental Biology and Medicine, 237(2), 167–177. 10.1258/ebm.2011.011321

Dudás, J., Dietl, W., Romani, A., Reinold, S., Glueckert, R., Schrott-Fischer, A., … Riechelmann, H. (2018). Nerve Growth Factor (NGF)—Receptor Survival Axis in Head and Neck Squamous Cell Carcinoma. International Journal of Molecular Sciences, 19(6), 1771. 10.3390/ijms19061771

Ekelem, C., Juhasz, M., Khera, P., & Mesinkovska, N. A. (2019). Utility of Naltrexone Treatment for Chronic Inflammatory Dermatologic Conditions. JAMA Dermatology, 155(2), 229–236. 10.1001/jamadermatol.2018.4093

Elmorsy, E. A., Saber, S., Hamad, R. S., Abdel-Reheim, M. A., El-kott, A. F., AlShehri, M. A., … Youssef, M. E. (2024). Advances in understanding cisplatin-induced toxicity: Molecular mechanisms and protective strategies. European Journal of Pharmaceutical Sciences, 203, 106939. 10.1016/j.ejps.2024.106939

Fanning, J., Hossler, C. A., Kesterson, J. P., Donahue, R. N., McLaughlin, P. J., & Zagon, I. S. (2012). Expression of the opioid growth factor–opioid growth factor receptor axis in human ovarian cancer. Gynecologic Oncology, 124(2), 319–324. 10.1016/j.ygyno.2011.10.024

France, C. P., Ahern, G. P., Averick, S., Disney, A., Enright, H. A., Esmaeli‐Azad, B., … Zapf, J. (2021). Countermeasures for Preventing and Treating Opioid Overdose. Clinical Pharmacology & Therapeutics, 109(3), 578–590. 10.1002/cpt.2098

Hamoui, M. Z., Rizvi, S., Arnouk, H., & Roberts, C. M. (2025). Putative Biomarkers for Prognosis, Epithelial-to-Mesenchymal Transition, and Drug Response in Cell Lines Representing Oral Squamous Cell Carcinoma Progression. Genes, 16(2), 209. 10.3390/genes16020209

Jiang, X.*, Wu, J.*, Wang, J., & Huang, R. (2019). Tobacco and oral squamous cell carcinoma: A review of carcinogenic pathways. Tobacco Induced Diseases, 17(April), 29. 10.18332/tid/105844

Li, Z., You, Y., Griffin, N., Feng, J., & Shan, F. (2018). Low-dose naltrexone (LDN): A promising treatment in immune-related diseases and cancer therapy. International Immunopharmacology, 61, 178–184. 10.1016/j.intimp.2018.05.020

Liu, W. M., Scott, K. A., Dennis, J. L., Kaminska, E., Levett, A. J., & Dalgleish, A. G. (2016). Naltrexone at low doses upregulates a unique gene expression not seen with normal doses: Implications for its use in cancer therapy. International Journal of Oncology, 49(2), 793–802. 10.3892/ijo.2016.3567

Liubchenko, K., Kordbacheh, K., Khajehdehi, N., Visnjevac, T., Ma, F., Khan, J. S., … Visnjevac, O. (2021). Naltrexone’s Impact on Cancer Progression and Mortality: A Systematic Review of Studies in Humans, Animal Models, and Cell Cultures. Advances in Therapy, 38(2), 904–924. 10.1007/s12325-020-01591-9

Ma, M., Wang, X., Liu, N., Shan, F., & Feng, Y. (2020). Low-dose naltrexone inhibits colorectal cancer progression and promotes apoptosis by increasing M1-type macrophages and activating the Bax/Bcl-2/caspase-3/PARP pathway. International Immunopharmacology, 83, 106388. 10.1016/j.intimp.2020.106388

McLaughlin, P J, Levin, R. J., & Zagon, I. S. (1999). Regulation of human head and neck squamous cell carcinoma growth in tissue culture by opioid growth factor. International Journal of Oncology, 14(5), 991–998. 10.3892/ijo.14.5.991

McLaughlin, Patricia J., Levin, R. J., & Zagon, I. S. (2003). Opioid growth factor (OGF) inhibits the progression of human squamous cell carcinoma of the head and neck transplanted into nude mice. Cancer Letters, 199(2), 209–217. 10.1016/s0304-3835(03)00341-0

McLaughlin, Patricia J., Stack, B. C., Levin, R. J., Fedok, F., & Zagon, I. S. (2003). Defects in the opioid growth factor receptor in human squamous cell carcinoma of the head and neck. Cancer, 97(7), 1701–1710. 10.1002/cncr.11237

McLaughlin, Patricia J, & Zagon, I. S. (2006). Progression of squamous cell carcinoma of the head and neck is associated with down-regulation of the opioid growth factor receptor. International Journal of Oncology, 28(6), 1577–1583.

McLaughlin, Patricia J., & Zagon, I. S. (2015). Duration of opioid receptor blockade determines biotherapeutic response. Biochemical Pharmacology, 97(3), 236–246. 10.1016/j.bcp.2015.06.016

McLaughlin, Patricia J, Zagon, I. S., Park, S. S., Conway, A., Donahue, R. N., & Goldenberg, D. (2009). Growth inhibition of thyroid follicular cell-derived cancers by the opioid growth factor (OGF) - opioid growth factor receptor (OGFr) axis. BMC Cancer, 9(1), 369. 10.1186/1471-2407-9-369

Minhas, S., Kashif, M., Altaf, W., Afzal, N., & Nagi, A. H. (2017). Concomitant-chemoradiotherapy-associated oral lesions in patients with oral squamous-cell carcinoma. Cancer Biology & Medicine, 14(2), 176–182. 10.20892/j.issn.2095-3941.2016.0096

Miskoff, J. A., & Chaudhri, M. (2018). Low Dose Naltrexone and Lung Cancer: A Case Report and Discussion. Cureus, 10(7), e2924. 10.7759/cureus.2924

Mokhtari, R. B., Homayouni, T. S., Baluch, N., Morgatskaya, E., Kumar, S., Das, B., & Yeger, H. (2017). Combination therapy in combating cancer. Oncotarget, 8(23), 38022–38043. 10.18632/oncotarget.16723

Montero, P. H., & Patel, S. G. (2015). Cancer of the Oral Cavity. Surgical Oncology Clinics of North America, 24(3), 491–508. 10.1016/j.soc.2015.03.006

Qu, N., Meng, Y., Handley, M. K., Wang, C., & Shan, F. (2021). Preclinical and clinical studies into the bioactivity of low-dose naltrexone (LDN) for oncotherapy. International Immunopharmacology, 96, 107714. 10.1016/j.intimp.2021.107714

Sano, D., Xie, T.-X., Ow, T. J., Zhao, M., Pickering, C. R., Zhou, G., … Myers, J. N. (2011). Disruptive TP53 Mutation Is Associated with Aggressive Disease Characteristics in an Orthotopic Murine Model of Oral Tongue Cancer. Clinical Cancer Research, 17(21), 6658–6670. 10.1158/1078-0432.ccr-11-0046

Seoane-Romero, J.-M., Vázquez-Mahía, I., Seoane, J., Varela-Centelles, P., Tomás, I., & López-Cedrún, J.-L. (2012). Factors related to late stage diagnosis of oral squamous cell carcinoma. Medicina Oral, Patología Oral y Cirugía Bucal, 17(1), e35–e40. 10.4317/medoral.17399

Shah, R., Shah, H., Thakkar, K., & Parikh, N. (2023). Conventional Therapies of Oral Cancers: Highlights on Chemotherapeutic Agents and Radiotherapy, Their Adverse Effects, and the Cost Burden of Conventional Therapies. Critical Reviews in Oncogenesis, 28(2), 1–10. 10.1615/critrevoncog.2023046835

Stockdale, D. P., Titunick, M. B., Biegler, J. M., Reed, J. L., Hartung, A. M., Wiemer, D. F., … Neighbors, J. D. (2017). Selective opioid growth factor receptor antagonists based on a stilbene isostere. Bioorganic & Medicinal Chemistry, 25(16), 4464–4474. 10.1016/j.bmc.2017.06.035

Tan, Y., Wang, Z., Xu, M., Li, B., Huang, Z., Qin, S., … Huang, C. (2023). Oral squamous cell carcinomas: state of the field and emerging directions. International Journal of Oral Science, 15(1), 44. 10.1038/s41368-023-00249-w

Toljan, K., & Vrooman, B. (2018). Low-Dose Naltrexone (LDN)—Review of Therapeutic Utilization. Medical Sciences, 6(4), 82. 10.3390/medsci6040082

Wang, X., Zhang, H., & Chen, X. (2019). Drug resistance and combating drug resistance in cancer. Cancer Drug Resistance, 2(2), 141–160. 10.20517/cdr.2019.10

Warnakulasuriya, S. (2009). Global epidemiology of oral and oropharyngeal cancer. Oral Oncology, 45(4–5), 309–316. 10.1016/j.oraloncology.2008.06.002

Zagon, I. S., Hytrek, S. D., & McLaughlin, P. J. (1996). Opioid growth factor tonically inhibits human colon cancer cell proliferation in tissue culture. American Journal of Physiology-Regulatory, Integrative and Comparative Physiology, 271(3), R511–R518. 10.1152/ajpregu.1996.271.3.r511

Zagon, Ian S. (1988). Endogenous opioid systems and neural cancer: Transmission and scanning electron microscopic studies of murine neuroblastoma in tissue culture. Brain Research Bulletin, 21(5), 777–784. 10.1016/0361-9230(88)90046-9

Zagon, Ian S., Donahue, R. N., & McLaughlin, P. J. (2009). Opioid growth factor-opioid growth factor receptor axis is a physiological determinant of cell proliferation in diverse human cancers. American Journal of Physiology-Regulatory, Integrative and Comparative Physiology, 297(4), R1154–R1161. 10.1152/ajpregu.00414.2009

Zagon, Ian S, & McLaughlin, P. J. (2006). Opioid growth factor receptor is unaltered with the progression of human pancreatic and colon cancers. International Journal of Oncology, 29(2), 489–494.

Zagon, Ian S, Porterfield, N. K., & McLaughlin, P. J. (2012). Opioid growth factor – opioid growth factor receptor axis inhibits proliferation of triple negative breast cancer. Experimental Biology and Medicine, 238(6), 589–599. 10.1177/1535370213489492

Zagon, Ian S., Verderame, M. F., & McLaughlin, P. J. (2003). The expression and function of the OGF–OGFr axis – a tonically active negative regulator of growth – in COS cells. Neuropeptides, 37(5), 290–297. 10.1016/j.npep.2003.07.001

Zagon, I.S., & McLaughlin, P. J. (1990). Opioid antagonist (naltrexone) stimulation of cell proliferation in human and animal neuroblastoma and human fibrosarcoma cells in culture. Neuroscience, 37(1), 223–226. 10.1016/0306-4522(90)90207-k

Zanoni, D. K., Montero, P. H., Migliacci, J. C., Shah, J. P., Wong, R. J., Ganly, I., & Patel, S. G. (2019). Survival outcomes after treatment of cancer of the oral cavity (1985–2015). Oral Oncology, 90, 115–121. 10.1016/j.oraloncology.2019.02.001

