## Supplementary Figures for "Naltrexone has variable and schedule-dependent effects on oral squamous cell carcinoma cells"

**Supplementary Figure S1**

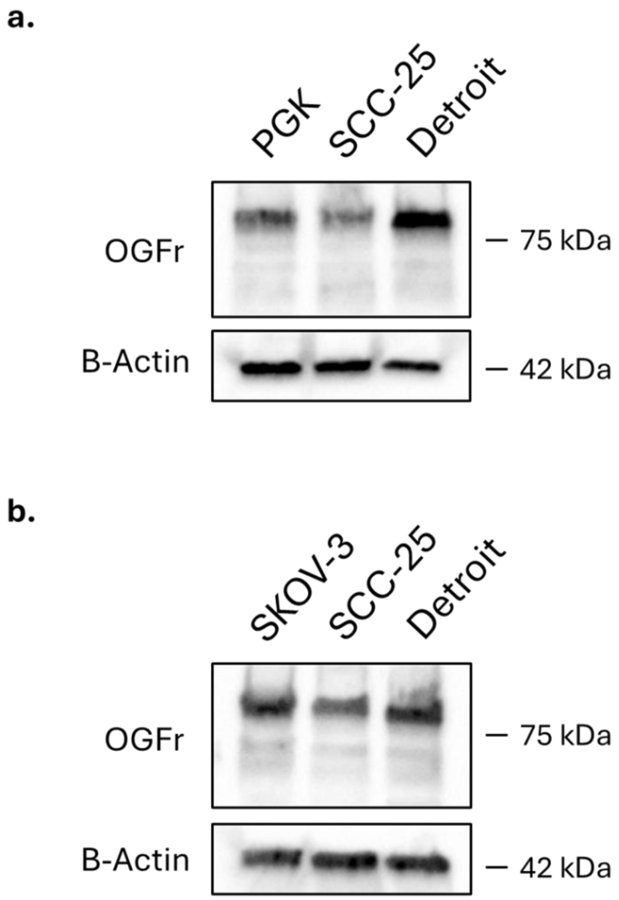


**Supplementary Figure S1. a.** Western blot showing relative expression of OGFr in OSCC cell lines and in normal control cells (PGK). Detroit 562 shows the highest expression. **b.** Western blot confirming similar levels of OGFr in SKOV-3 ovarian cancer cells as in OSCC lines. n=1 for each.

**Supplementary Figure S2**

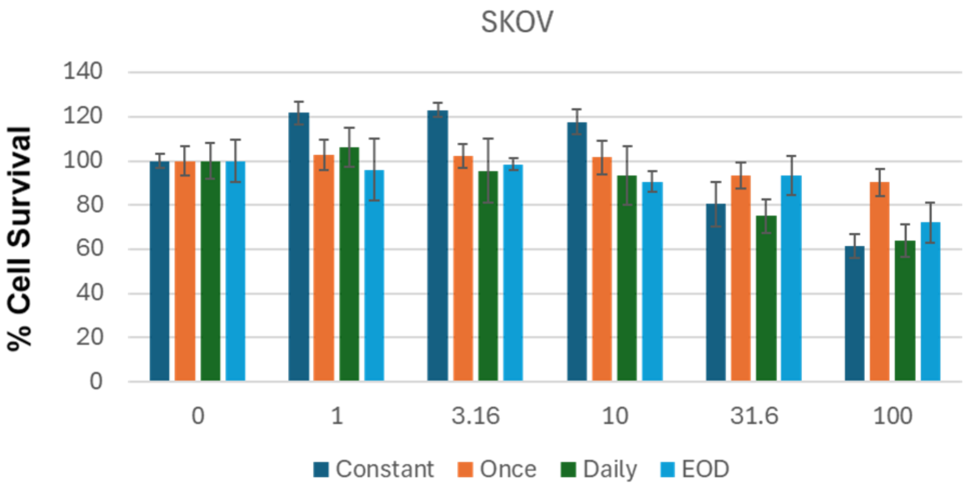


**Supplementary Figure S2.** Single experiment with expanded dose range shows NTX toxicity in SKOV-3 cells at high doses independent of dosing schedule. Graph shows mean ± SD of technical replicates. X-axis, dose of NTX in μM.
